# Past and present caregiving experiences impact prefrontal connectivity and recall for attachment-schema narratives

**DOI:** 10.1101/2024.09.13.612953

**Authors:** Caroline S. Lee, Samantha S. Cohen, Samuel Hutchinson, Nim Tottenham, Christopher Baldassano

## Abstract

We investigated how past and current caregiving experiences impacted emotional event processing by examining inter-subject functional correlation in 7- to 15-year-olds during narrative movies depicting separation and reunion. Early adversity impacted amygdala interactions with the hippocampus, but medial prefrontal-amygdala connectivity and the content of later recall was shaped by attachment security to current caregivers. This suggests that current attachment can have a redemptive effect on attachment schemas formed in early childhood.

---

Early caregiving experiences can play a major role in how children process and emotionally respond to interpersonal interactions throughout their lives. Recent reviews of the neural and behavioral impacts of caregiving-related early childhood adversities (crEAs), including caregiving instability, abuse, and neglect, have proposed that these experiences specifically disrupt the construction of *affective schemas*: semanticized knowledge and predictions about how socio-emotional events are likely to unfold (Tottenham, 2020; Tottenham and Vannucci, 2025). This internal model of emotional expectations (about, for example, whether a caregiver will be responsive to the child’s needs) can then impact how new experiences are experienced and remembered (Bowlby, 1969; Bretherton, 1992). Based on work using general event scripts (Baldassano et al., 2018; Kurby & Zacks, 2008; Gilboa & Marlatte, 2017), we hypothesized that disordered affective schemas could impede the typical top-down regulation of emotional signals in amygdala by medial prefrontal cortex (mPFC). A key question is whether these prefrontal-amygdala interactions are most related to caregiving stability in *early* life (when attachment schemas are initially constructed) or to attachment security with *current* caregivers (indicating that these schemas can be updated in later childhood).

Rather than using static or highly-simplified stimuli to evoke an attachment schema, we presented children with one of two short naturalistic movies that depicted a separation and reunion narrative and measured the stimulus-driven connectivity of the amygdala using inter-subject functional correlation (ISFC; Simony et al, 2016). Children also freely recalled the events of the video they watched, allowing us to test whether early instability or current attachment impacted the schematic content of narrative recall.

Participants (N = 187; ages 7 – 15) included children with a variety of caregiver experiences, assessed in two ways (Figure 1A): (1) the stability of their early life caregiver environment was characterized as either “ stable” (N=78 for fMRI analyses, N=104 for recall analyses) or “ unstable” (N=51, N=82), based on whether they had ever experienced a change in their primary caregivers (e.g. being removed from the care of their parents or moving between foster families) and (2) the attachment relationship quality with their current caregiver was characterized as “ strong” (N=54, N=87) or “ weak” (N=48, N=79). Brain activity was measured while each participant viewed a brief movie clip depicting an attachment narrative, after which each participant verbally recalled the movie to an experimenter (Figure 1B). Two different narrative movies were used, constructed from scenes from *Homeward Bound* or *The Little Princess* in which a character is (1) initially together with their caregiver; (2) separated from their caregiver; (3) searching for their caregiver; and (4) reunited with their caregiver (Figure 1C).

**Figure 1:**
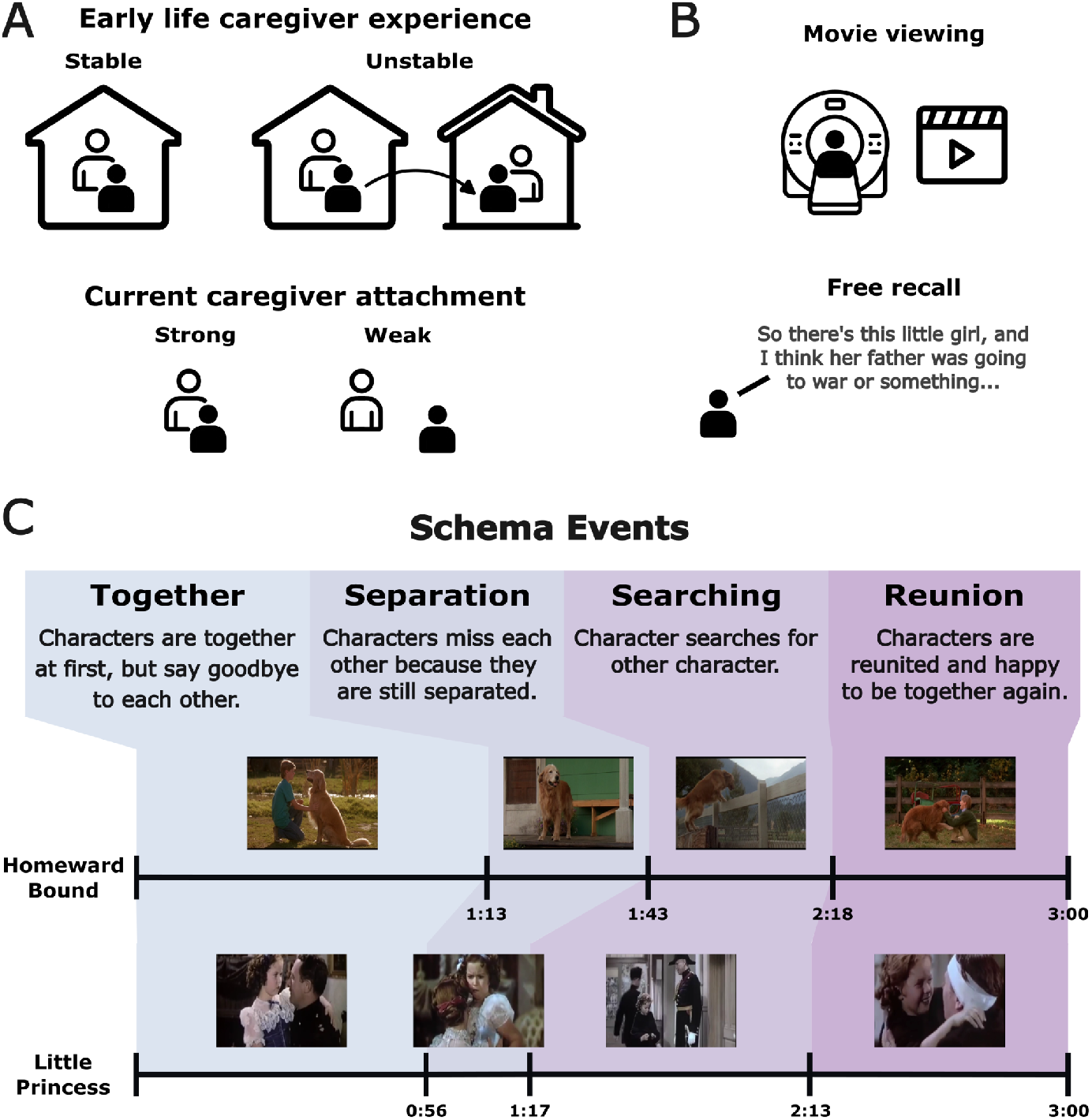
Study population and design. **A**. Children either had a single caregiving environment (stable) or experienced a permanent change in their caregivers (unstable) in early childhood. Separately, we measured the strength of each child’s attachment to their current caregivers, creating Strong attachment and Weak attachment groups. **B**. Children watched a three-minute movie clip during an fMRI session (top) and were later asked to recall what they remembered from the movie to an experimenter (bottom) outside of the scanner. **C**. Each movie clip (“ Homeward Bound” or “ Little Princess” ) was edited to depict the same sequence of four events associated with an attachment schema (“ Together” , “ Separation” , “ Searching” , “ Reunion” ).

We measured the univariate timecourse (in each hemisphere) of activity in the amygdala, 100 neocortical parcels, and hippocampus. For children with each kind of attachment experience (e.g. those with stable early caregivers) who viewed the same attachment narrative, we computed the stimulus-driven ISFC connectivity for each brain ROI and the amygdala as the correlation between the average amygdala response in one half of the group and an average ROI response in the other half (Figure 2A). Measured ISFC values were averaged across the two movies and then statistically compared between subgroups (stable versus unstable early caregiving or strong versus weak current attachment).

**Figure 2:**
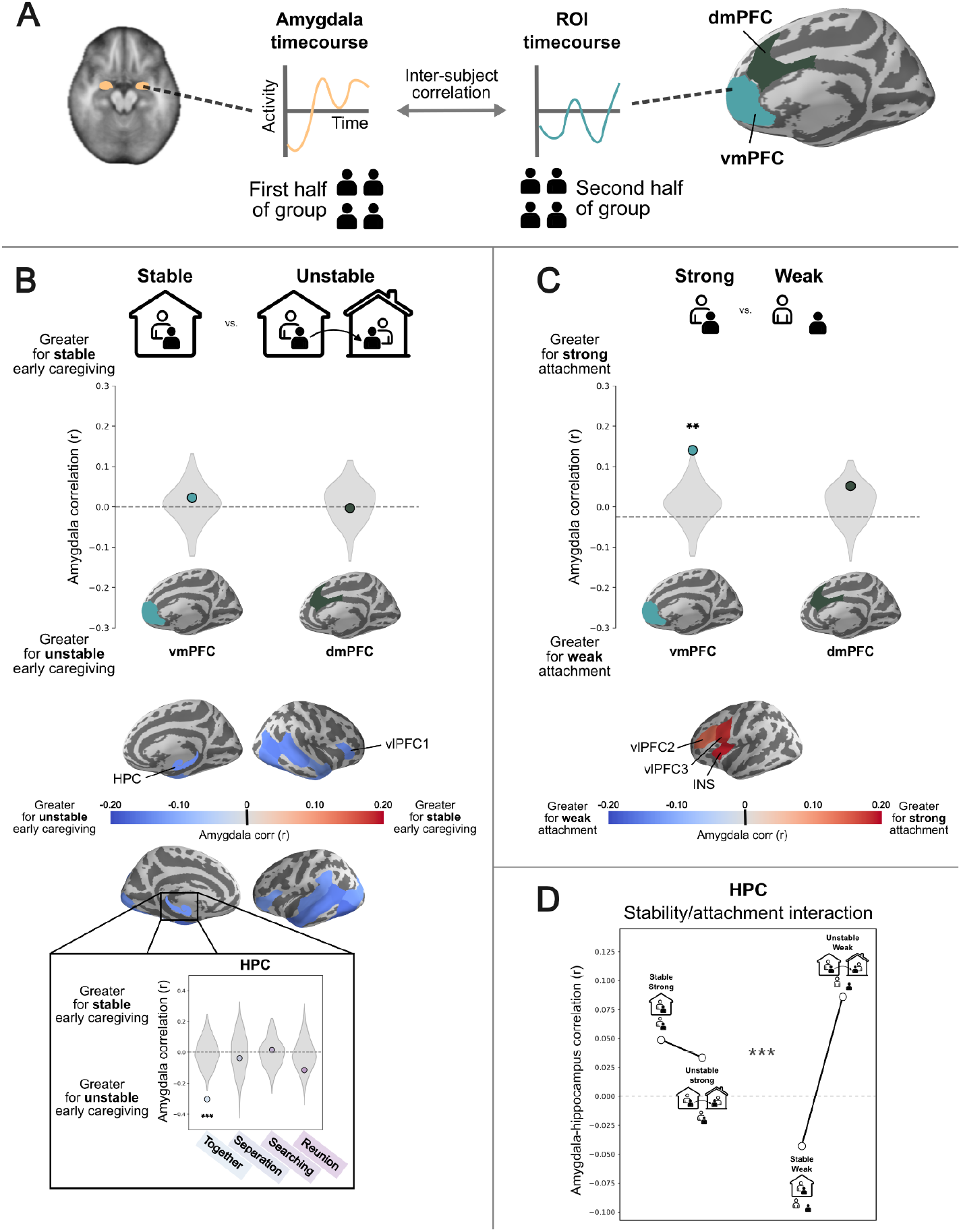
Inter-subject functional correlation (ISFC) for attachment narratives between the amygdala and other brain regions. **A**. To compute ISFC, a group of participants (e.g. those with stable caregiving environments) is randomly divided in half. The mean amygdala response timecourse is computed in one half, and the mean response timecourse of another ROI (such as vmPFC or dmPFC) is computed in the other half. The ISFC is the correlation between these two timecourses, reflecting the degree of shared stimulus-driven information between the amygdala and this ROI for this group. **B**. Caregiving environments in early childhood were not significantly related to amygdala-mPFC coupling, but a whole-brain analysis found that unstable caregiving was associated with stronger amygdala coupling to sensory regions in visual and auditory cortex, as well as the hippocampus (HPC) and ventrolateral PFC parcel (vlPFC1). (Inset) The HPC effects were driven by the beginning of the narratives, during the initial event (Together), for children who experienced unstable caregiving environments. **C**. Examining medial PFC parcels implicated in previous research on affective schemas, we found greater amygdala coupling to ventromedial PFC in children with strong levels of current attachment. Strong current attachment was also associated with greater amygdala coupling to ventrolateral prefrontal cortex (vlPFC2, vlFPC3) and the insula (INS) in the left hemisphere. **D**. Among prefrontal and hippocampal regions showing effects of stability or attachment, there was an interaction effect only in the hippocampus. The increased amygdala connectivity associated with unstable caregiving was only present among children that had weak attachment to their current caregivers. Brain maps are thresholded at q<0.05. * p < 0.05, ** p < 0.01, *** p < 0.001

Both early life caregiver experience and current feelings of attachment security had a significant association with amygdala connectivity patterns, tested in *a priori* ventromedial PFC (vmPFC) and dorsomedial PFC (dmPFC) ROIs and in a whole-brain parcellation of the neocortex and hippocampus. In children with unstable early caregiving experiences, the amygdala did not significantly differ in its connectivity to mPFC, but was more connected to sensory regions in the auditory and visual cortex, the hippocampus, and a bilateral ventrolateral PFC parcel (Figure 2B). We conducted an exploratory analysis to measure ISFC within each of the four schematic events in the movies, and found that this effect was significantly driven only by event 1 in the hippocampus (p = 0.0004; corrected for multiple comparisons), suggesting that the the initial presentation of a caregiving relationship triggers interactions with episodic memory for children with an unstable caregiving history. Children with stronger current feelings of attachment security, in contrast, exhibited greater amygdala coupling in vmPFC (p = 0.01), as well as in vlPFC regions and the anterior insula (Figure 2C). In all regions, this enhanced connectivity was distributed throughout the movie and was not significantly driven by any specific event.

Additionally, we tested for interactions between caregiving stability and current attachment on amygdala connectivity in the mPFC, vlPFC, insula, and hippocampus parcels identified in the previous analyses (Figure 2D). We found a significant interaction in the hippocampus (p = 0.0006; corrected for multiple comparisons); unstable caregiving increased hippocampal-amygdala connectivity only among children with weak current attachment security, suggesting that a strong attachment to current caregivers may mitigate disruptions caused by unstable early caregiving.

We next examined the content of each child’s recall of the narrative movies by applying a transformer-based language model (Devlin et al., 2018) to the recall transcripts. This model converts each recall into a vector, such that measuring the cosine similarity between vectors provides a quantitative measure of the semantic similarity between recalls (Figure 3A). We found that the model similarities were related to neural activity (Figure 3B); controlling for age, verbal IQ, and gender, the semantic similarity between a pair of participants was significantly predicted by their similarity in neural response timecourses in HPC (p = 0.02) and vlPFC2 (p = 0.007), and marginally in vlPFC3 (p = 0.06; all p values corrected for multiple comparisons).

**Figure 3:**
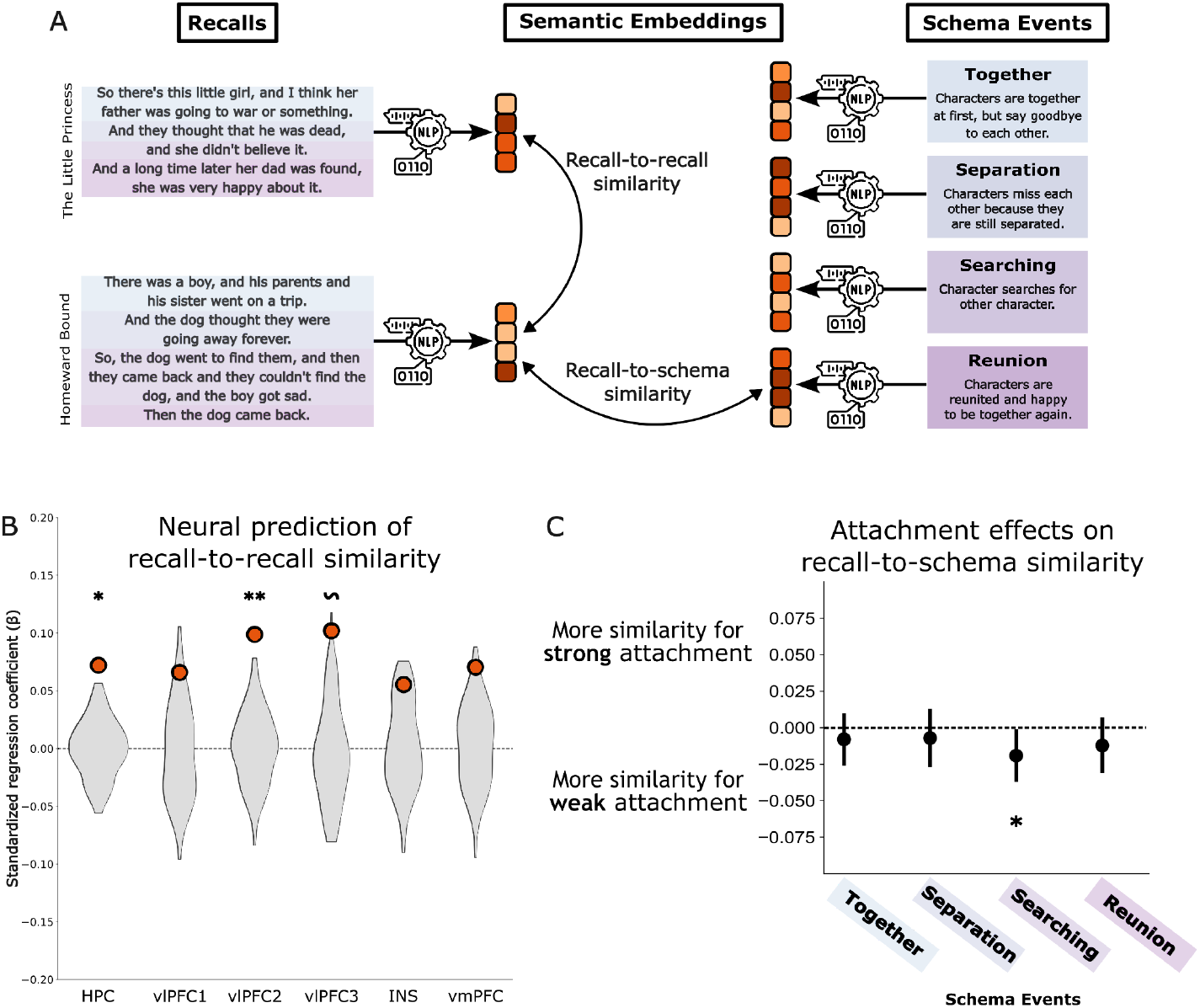
Analysis of semantic content of participant recalls. **A**. Vector embeddings of each participant’s recall and each event in the attachment schema were generated using a sentence transformer model. The cosine similarity between these vectors provides a measure of semantic similarity, either between recalls from different participants or between a recall and schematic event template. **B**. For HPC, vlPFC2, and vmPFC, participants with more similar neural responses also had recalls with more similar semantic content. **C**. The recalls of children with weaker current caregiver attachment had semantic content that was more influenced by the Searching event, when the characters in the story are apart from each other and reunion is uncertain. * p < 0.05, ** p < 0.01, ∼ p <0.06

To test whether early caregiving or current attachment influenced the schematic content of recalls, we also applied the language model to a text description of each of the four stages of our attachment script and measured the semantic similarity between each recall and each schematic event. Controlling for age, verbal IQ, and gender, we found that children with weaker attachment security had recalls with significantly more semantic content related to the Searching phase of the attachment script (p = 0.04) (Figure 3B), suggesting that weak attachment security biases memories of caregiving experiences to produce an overrepresentation of events in which the characters are separated. There was no significant effect on the schematic content of recall associated with early caregiving stability.

In summary, we found that early life caregiving and weaker current caregiver attachment altered both functional connectivity and memory recall in response to a naturalistic attachment video. In the brain, the way in which the amygdala interacts with the neocortex and hippocampus showed a marked dissociation between past experience and current attachment. Medial prefrontal connectivity to the amygdala was disrupted specifically in children with low levels of attachment security rather than in all children who experienced crEAs, suggesting some degree of plasticity for affective schemas based on more recent experiences. Low attachment security also impacted amygdala connectivity with the insula, which has been shown to exhibit predictive signals for narratives (Lee, Aly, & Baldassano, 2021), suggesting that children with secure attachments could use schema-based predictions (i.e. that the caregiver will return after an absence) to modulate amygdala responses. Attachment security also impacted later recall, as children with secure attachments focused less on the aspects of the narrative involving the character searching for a caregiver.

Early instability did have an effect on other components of amygdala connectivity, predominantly increasing interactions with sensory and episodic memory systems. Stronger connectivity to sensory input is consistent with a tradeoff between top-down and bottom-up modulation of the amygdala, with greater amygdala responsivity to the current sensory input in the absence of a well-organized internal schema. Instability increased connectivity between the amygdala and the hippocampus, as in animal models of chronic stress (Ghosh et al., 2013), but only among children with weak current attachment. This suggests that strong attachments to caregivers can serve a protective function in children with a history of crEA exposure.

## Online Methods

### Participants

222 children (7 - 15 years) were recruited as a part of a larger longitudinal study on disruptions in caregiving (see Fields et al., 2021 and Nikolaidis et al., 2022 for recruitment details). 135 of the cohort experienced disruptions in caregiving (e.g., institutionalization, domestic and international foster care, temporary/permanent separations from parents) and a comparison group of children did not (N = 87). All recruitment and data collection procedures were approved by the Columbia University IRB.

Caregiving instability was characterized by whether or not a child had a switch in primary caregiver or not, regardless of group (disrupted or comparison). This was assessed from a verbally reported timeline of each child’s lifetime caregiving placements from their current legal guardian. A caregiving switch was characterized as a separation from biological parents (including those at, or shortly after birth to account for switches between the pre- and post-natal environment), orphanage placements, foster care placements, kinship care placements, or moves from one biological parent’s home to another (on-going co-parenting arrangements did not count).

Attachment security was assessed from the 30-item Attachment strategies questionnaire (Kerns et al., 1996). Based on prior work, children with scores below 3.0 were defined as the “ weak” security group, while those with scores above 3.27 made up the “ strong” security group. 7 participants had incomplete phenotype data (e.g. missing attachment security scores) and were dropped from fMRI and behavioral analyses.

Both groups, current attachment security and caregiving stability, were further examined for over-representation in age, gender, and verbal proficiency. First, none of the attachment security levels were significantly over-represented in either of the caregiving stability groups (Cramer’s V = 0.13). Neither gender, male or female, were over-represented in the caregiving stability groups (Cramer’s V = 0.04) or the attachment security levels (Cramer’s V = 0.03). Next, a one-way ANOVA was conducted to confirm that there were no significant effects of age in either of the caregiving stability groups [F(1, 98) = 0.60, p = 0.44] or in the attachment security groups [F(2, 97) = 0.18, p = 0.84]. And lastly, the effects of verbal proficiency in both groups were examined with a one-way ANOVA, with no significant differences between attachment security groups [F(2, 97) = 1.63, p = 0.20]. However, there was a significant difference in verbal proficiency between the two caregiving stability groups [F(1, 95) = 6.84, p = 0.01]. Regardless of significance, we add all factors (age, gender, and verbal proficiency) as regressors in our linear regression models to evaluate recall transcripts.

### Stimuli

Children saw one of two video stimuli, each three minutes in duration: a compilation from The Little Princess (1939) or a compilation from Homeward Bound (1993). Each stimulus was chosen and carefully edited to contain four scenes describing different fundamental themes of attachment: (1) Together, (2) Separation, (3) Searching, and (4) Reunion.

### MRI scanning protocol and preprocessing

Prior to scanning all participants completed a mock scanning session to acclimate to the scanner environment and practice lying still for data collection. Data was collected on Siemens Magnetom Prisma 3T MRI scanner using a 64-channel head coil. A whole-brain high-resolution T1-weighted anatomical scan (MPRAGE; 256 × 256 in-plane resolution; 256 mm FOV; 176 × 1 mm sagittal slices) was acquired for each participant for registration of functional data. Children were randomly assigned to watch one of the video stimuli which were presented via an MR-compatible screen and mirror attached to the head coil. The MRI data included in this manuscript were preprocessed using fMRIprep 20.2.1 (Esteban et al., 2018; Esteban, Blair, et al., 2018).

### Free recall protocol

After participants had completed their MR scan, an experimenter asked them to recall as much as they could from the video stimulus that they had viewed in the scanner. Recalls were obtained about 15 minutes after scan sessions on average. Their recall was recorded and transcribed. 186 total transcripts were included in the stimulus, excluding 36 participants who either did not participate in the recall portion of the experiment or had transcripts without scorable content.

### Anatomical and functional data preprocessing

The MRI data results included in this manuscript come from preprocessing performed using fMRIPprep 20.2.1 (Esteban et al., 2018; Esteban, Blair, et al., 2018; RRID:SCR_016216), based on Nipype 1.5.1 (Gorgolewski et al., 2011; Gorgolewski et al., 2018; RRID:SCR_002502). A total of two T1-weighted (T1w) images were present for the majority of the dataset. T1w images were corrected for intensity non-uniformity (INU) using ‘N4BiasFieldCorrection’ (Tustison et al., 2010), distributed with ANTS 2.3.3 (Avants et al., 2008, RRID:SCR_004757). The T1w-reference was then skull-stripped using ‘antsBrainExtraction.sh’ workflow (from ANTs), using OASIS30ANTs as a target template. Brain tissue segmentation of cerebrospinal fluid (CSF), white-matter (WM) and gray-matter (GM) was performed on the brain-extracted T1w using fast (FSL 5.0.9, RRID:SCR_002823, Zhang, Brady, and Smith 2001). A T1w-reference map was computed after registration of T1w images (after INU-correction) using mri_robust_template (FreeSurfer 6.0.1, Reuter, Rosas, and Fischl; 2010). Brain surfaces were reconstructed using recon-all (FreeSurfer 6.0.1, RRID:SCR_001847, Dale, Fischl, and Sereno 1999), and the brain mask estimated previously was refined with a custom variation of the method to reconcile ANTs-derived and FreeSurfer-derived segmentations of the cortical gray-matter of Mindboggle (RRID:SCR_002438, Klein et al. 2017).

The amygdala was defined using an MNI template in standard space (MNI152NLin2009cAsym). Nonlinear registration to this template was performed with antsRegistration (ANTs 2.3.3) using brain-extracted versions of both the T1w reference and the T1w template. The following template was selected for spatial normalization: ICBM 152 Nonlinear Asymmetrical template version 2009c (Fonov et al., 2009), RRID:SCR_008796; TemplateFlow ID: MNI152NLin2009cAsym).

### Functional data preprocessing

First, a reference volume and its skull-stripped version were generated using the custom methodology of fMRIPrep. A B0-nonuniformity map (or fieldmap) was estimated based on two echo-planar imaging (EPI) references with opposing phase-encoding directions, with 3dQwarp Cox and Hyde (1997) (AFNI 20160207). Based on the estimated susceptibility distortion, a corrected EPI (echo-planar imaging) reference was calculated for a more accurate co-registration with the anatomical reference. The BOLD reference was then co-registered to the T1w reference using bbregister (FreeSurfer) which implements boundary-based registration (Greve and Fischl 2009). Co-registration was configured with six degrees of freedom. Head-motion parameters with respect to the BOLD reference (transformation matrices, and six corresponding rotation and translation parameters) are estimated before any spatiotemporal filtering using mcflirt (FSL 5.0.9, Jenkinson et al. 2002). The BOLD time-series were resampled onto fsaverage6 for analyses on the cortical surface and MNI152NLin2009cAsym space for analysis of the amygdala. The BOLD time-series were resampled onto their original, native space by applying a single, composite transform to correct for head-motion and susceptibility distortions. These resampled BOLD time-series will be referred to as preprocessed BOLD.

Several confounding time-series were calculated based on the preprocessed BOLD: framewise displacement (FD) and three region-wise global signals. FD was computed using two formulations following Power (absolute sum of relative motions, Power et al. (2014)) and Jenkinson (relative root mean square displacement between affines, Jenkinson et al. (2002)). The DVARS variable was derived from the derivative of the estimated relative (frame-to-frame) bulk head motion calculated using the root mean squared approach of Jenkinson et al. (2002). FD is calculated for each functional run using the implementation in Nipype (following the definitions by Power et al. 2014). The three global signals are extracted within the CSF, the WM, and the whole-brain masks. The confound time series derived from head motion estimates and global signals were expanded with the inclusion of temporal derivatives and quadratic terms for each (Satterthwaite et al. 2013). Frames that exceeded a threshold of 0.5 mm FD or 1.5 standardized DVARS were annotated as motion outliers. Of the 179 participants with complete fMRI data, of which 7 were dropped due to incomplete phenotype data, the number of motion outliers did not significantly differ in either split (caregiving stability: t(154) = -0.42, p = 0.68; attachment security: t(141) = 0.47, p = 0.64). Participants whose timeseries contained more than 45 TRs (20% of all TRs) of motion outliers were excluded from further analyses due to excessive motion. Discrete cosine filters were extracted with 125 s cut-off. The nuisance regressors, as well as the head-motion estimates, were placed in a confounds file and subsequently regressed out of the BOLD time series separately for the cortex and the amygdala using custom python scripts. All resamplings can be performed with a single interpolation step by composing all the pertinent transformations (i.e. head-motion transform matrices, susceptibility distortion correction when available, and co-registrations to anatomical and output spaces). Gridded (volumetric) resamplings were performed using antsApplyTransforms (ANTs), configured with Lanczos interpolation to minimize the smoothing effects of other kernels (Lanczos 1964). Non-gridded (surface) resamplings were performed using mri_vol2surf (FreeSurfer).

Many internal operations of fMRIPrep use Nilearn 0.6.2 (Abraham et al. 2014, RRID:SCR_001362), mostly within the functional processing workflow. For more details of the pipeline, see the section corresponding to workflows in fMRIPrep’s documentation. After these preprocessing steps were taken, the cortical surface was parcellated into the 100 parcel seven network parcellation from Schaefer et al., 2018.

### Inter-subject functional correlation (ISFC) analysis

After the fMRI data was preprocessed and the cortical data was parcelated on the cortical surface (Schaefer et al., 2018) subjects were divided based on the number of caregiver switches that they had experienced and their current attachment security. The following ISFC analyses were run within two distinct groupings of the data: those who had experienced caregiving switches vs. those who had not, and those who felt a high level of attachment security with their current caregiver vs. those who felt a low level of attachment security with their current caregiver.

An ISFC analysis was run within each group, following previous research (Simony et al., 2016). For this analysis, we followed a protocol to measure inter-subject correlation established previously (Cohen, Tottenham, & Baldassano, 2022) wherein each group was randomly split into two subgroups ten times. In one subgroup all of their responses from one cortical parcel were averaged, and in the other subgroup, all of their responses from one hemisphere’s amygdala were averaged. These group-averaged responses were then correlated to establish an ISFC value between each parcel and one hemisphere’s amygdala. As we did not expect hemispheric differences in amygdala connectivity, the ISFC values for each cortical parcel were averaged across the amygdala from each hemisphere. Similarly, we then averaged ISFC values across both hemispheres of each ROI. To compare the ISFC values between groups (e.g. caregiver switches vs. no caregiver switches) we subtracted the ISFC values for one group from those of the other for all cortical parcels. All of these analyses were run separately for subjects that had watched one of the two video stimuli, and then averaged across stimuli.

ISFC values for event-specific timecourses were computed in a similar fashion as the full timecourse, but with responses from each individual’s amygdala and ROI taken only from a selection of consecutive TRs belonging to each respective event (Event 1: 0 - 70 TRs and 0 - 91 TRs in The Little Princess and Homeward Bound, respectively; Event 2: 71 - 96 and 92 - 129; Event 3: 97 - 166 and 130 - 173; and Event 4: 167 - 225 and 174-225). All group and hemisphere averaging across ISFC values were performed as for the full-timecourse analysis described above.

Interaction effects of attachment and caregiving were examined by dividing participants into four groups, based on a 2×2 split according to both stable vs unstable caregiving environments and strong vs weak attachment. ISFC was computed within each of the four groups. The magnitude of the interaction effect was defined as the root mean square residual of each group’s ISFC after taking into account both the main effects of attachment and caregiving. All averaging across groups, amygdala hemispheres, and ROI hemispheres were consistent with the main analysis.

Statistical significance for all ISFC analyses was assessed via permutation tests, in which the analyses were run 100 times for groups in which the life experience and attachment variables of interest were randomly shuffled across subjects. The null distribution was modeled as a Normal distribution, and a p value was computed from the survival function for this distribution at the real value from the original analysis (measuring the fraction of the null distribution in which the effect exceeds the observed effect).

### Free recall analysis

After removing stop-words from recall transcripts, we extracted an embedding from each transcript using a transformer-based language representation model (Sentence BERT; Devlin et al., 2018). We conducted a representational similarity analysis between participants examining the degree to which neural activity can predict remembered semantic content. Pairwise correlations between the neural response timecourses of participants were z-scored and then used in a linear regression model to predict the cosine similarities of recall embeddings in corresponding participant pairs. A linear models were fit to neural activity from each vlPFC parcel, the insula, and bilateral hippocampus, along with nuisance regressors of pairwise age differences, verbal proficiency (IQ), and sex differences. Statistical significance for the regression coefficient for neural activity was then computed by shuffling neural activity across participants prior to generating pairwise correlations, resulting in 100 betas in each null distribution. As above, this null distribution was approximated as a Normal distribution to produce a p-value, which was then corrected for multiple comparisons across 5 parcels.

We also measured the proximity between each child’s transcribed recall and the attachment narrative. We created four schematic event templates abstractly describing each of the four events and used Sentence BERT to convert them to embeddings. We then generated 4 proximity scores per participant by taking the cosine distance from recall embeddings to each event’s embeddings and subtracting the distance from 1, where larger values denote more proximity between the event and recall. Similar to the ISFC analysis, we examined proximity scores between the highest versus lowest attachment groups and between children with stable versus unstable caregiving histories. We fit a linear regression model to each of the 4 events’ proximity scores, adding participants’ age, verbal IQ, and gender as regressors.

